# Topographic specificity of alpha power during auditory spatial attention

**DOI:** 10.1101/681361

**Authors:** Yuqi Deng, Inyong Choi, Barbara Shinn-Cunningham

**Affiliations:** Department of Biomedical Engineering, Boston University, Boston, MA, USA, 02215; Department of Communication Sciences and Disorders, University of Iowa, Iowa City, IA, 52242; Carnegie Mellon Neuroscience Institute, Carnegie Mellon University, Pittsburgh, PA 15213

**Keywords:** electroencephalography, selective attention, human behavior, parietal cortex

## Abstract

Visual and somatosensory spatial attention both induce parietal alpha (7-14 Hz) oscillations whose topographical distribution depends on the direction of spatial attentional focus. In the auditory domain, contrasts of parietal alpha power for leftward and rightward attention reveal a qualitatively similar lateralization; however, it is not clear whether alpha lateralization changes monotonically with the direction of auditory attention as it does for visual spatial attention. In addition, most previous studies of alpha oscillation did not consider subject-specific differences in alpha frequency, but simply analyzed power in a fixed spectral band. Here, we recorded electroencephalography in human subjects when they directed attention to one of five azimuthal locations. After a cue indicating the direction of an upcoming target sequence of spoken syllables (yet before the target began), alpha power changed in a task specific manner. Subject-specific peak alpha frequencies differed consistently between frontocentral electrodes and parieto-occipital electrodes, suggesting multiple neural generators of task-related alpha. Parieto-occipital alpha increased over the hemisphere ipsilateral to attentional focus compared to the contralateral hemisphere, and changed systematically as the direction of attention shifted from far left to far right. These results showing that parietal alpha lateralization changes smoothly with the direction of auditory attention as in visual spatial attention provide further support to the growing evidence that the frontoparietal attention network is supramodal.

## 1 Introduction

Visual spatial attention engages a well-studied frontoparietal network (e.g., Capotosto et al., 2009; Corbetta, 1998; He et al., 2007; Shulman et al., 2010). This network involves distinct regions in lateral frontal cortex that are separated by areas biased towards processing auditory inputs (Michalka et al., 2016; Noyce et al., 2017). The visual attention network also includes a series of retinotopic maps that start near primary visual sensory cortex and ascend along intraparietal sulcus (IPS; e.g., see Sereno et al., 2001; Swisher et al., 2007). Activity in these retinotopic maps, which represent contralateral space, is modulated by visual spatial attention; indeed, spatial attention alone can lead to activation in these areas, even in the absence of visual stimulation (Saygin and Sereno, 2008; Silver et al., 2005).

Many have argued that the frontoparietal network is supramodal, involved not just in visual spatial attention, but also in somatosensory and auditory spatial attention. Recent fMRI evidence supports this view. Specifically, auditory tasks involving spatial attention and spatial working memory engage the same lateral frontal cortex regions active during visual tasks (Michalka et al., 2016; Noyce et al., 2017). Auditory spatial tasks, but not non-spatial tasks, engage IPS (e.g., Alain et al., 2001; Arnott et al., 2004), although this activation seems to be restricted to later, higher-order maps without engaging the earlier IPS maps nearer to visual cortex (Michalka et al., 2016).

Neuroelectric imaging studies (using electro- and magnetoencephalography—EEG and MEG) reveal a strong signature of the direction of visual spatial attention, attributed to activity in the retinotopic IPS regions. When attention is directed to one side of space, there is typically an increase in neural oscillation power in the alpha range (7-14 Hz) from ipsilateral parietal cortex, and a decrease in alpha power from contralateral parietal cortex (Kelly et al., 2006; Thut et al., 2006; Worden et al., 2000; Wöstmann et al., 2016). This lateralization of parietal alpha power varies smoothly as the direction of visual spatial attention shifts, providing a readout of the direction of visual attentional focus (Foster et al., 2016; Rihs et al., 2007; Samaha et al., 2016; Worden et al., 2000). Given that parietal lobes primarily encode information about events that are in contralateral exocentric space, parietal alpha lateralization is thought to reflect a suppression of information (Foxe and Snyder, 2011; Klimesch, 2012; Klimesch et al., 2007; Romei et al., 2010). Specifically, in the parietal lobe ipsilateral to the direction of attention, alpha increases to suppress objects that are to be ignored, while in the parietal lobe contralateral to the direction of attention, alpha decreases to allow processing of an attended object (Ikkai et al., 2016).

A few studies have contrasted parietal alpha lateralization when *auditory* spatial attention is directed to the left versus to the right, and found a pattern that is qualitatively similar to that seen in visual spatial attention (Klatt et al., 2018; Tune et al., 2018). Yet, there is little known about whether auditory spatial attention varies monotonically as attentional focus shifts, as it does in vision (Rihs et al., 2007; Samaha et al., 2015; van Gerven and Jensen, 2009; Worden et al., 2000).

To study the effects of spatial auditory attention on alpha activity, we designed an auditory attention task in which listeners were cued at the start of each trial as to which of five spatial locations (varying in lateral position) would contain a target sequence. We measured EEG as while listeners were actively engaged in the auditory spatial attention task. We investigated how the alpha peak frequency varied across the scalp, and how each subject’s individualized parietal alpha frequency power distribution was modulated by the direction of attention. One challenge in studying alpha is that there may be significant differences across subjects in the alpha peak frequency, as well as multiple generators of alpha, which also might vary in their peak frequency as well as their topography on the scalp (Haegens et al., 2014). To enhance the sensitivity of our analysis, we therefore determined subject-specific estimates of parietal alpha frequency and analyzed how a narrow, 2 Hz wide band of power centered on this peak was modulated by the direction of attention (as opposed to analyzing the average power over the range of observed alpha, e.g., 7-14 Hz).

## 2 Materials and Methods

### 2.1 Participants

Thirty subjects (14 females, 18-30 years of age) participated in this study. All subjects had normal hearing (hearing thresholds better than 20dB at pure tone frequencies between 250 Hz and 8 kHz). All gave informed consent as approved by the Boston University Institutional Review Board. Two subjects were excluded from the study due to an inability to perform the task (percentage of correct responses equaled chance level). Subjects were asked to fill out an Edinburgh handedness inventory questionnaire (Oldfield, 1971) to determine their handedness preference. Fourteen out of the 28 remaining subjects were right-handed while the rest were left-handed.

### 2.2 Paradigm

Participants performed a spatial attention task in which they had to identify a target sequence of three spoken syllables from one direction while ignoring a distractor sequence of three similar syllables from another direction (Figure 1). At the start of each trial, a visual fixation dot appeared on the screen. Two seconds later, an auditory cue was played from one of five possible locations to indicate the spatial location of the upcoming target sequence. The target sequence and distractor sequence onsets were separated by 200 ms, allowing the neural responses elicited by the onsets of syllables in each stream to be temporally resolved. Within each stream, the syllable onsets were separated by 500 ms. To make sure that listeners engaged spatial attention (rather than being able to rely on temporal expectations), on half of the trials, the target stream began before the distractor, while in the other trials the distractor began first. The first (target or distractor) sequence began to play two seconds after the auditory cue.

**Figure 1.**
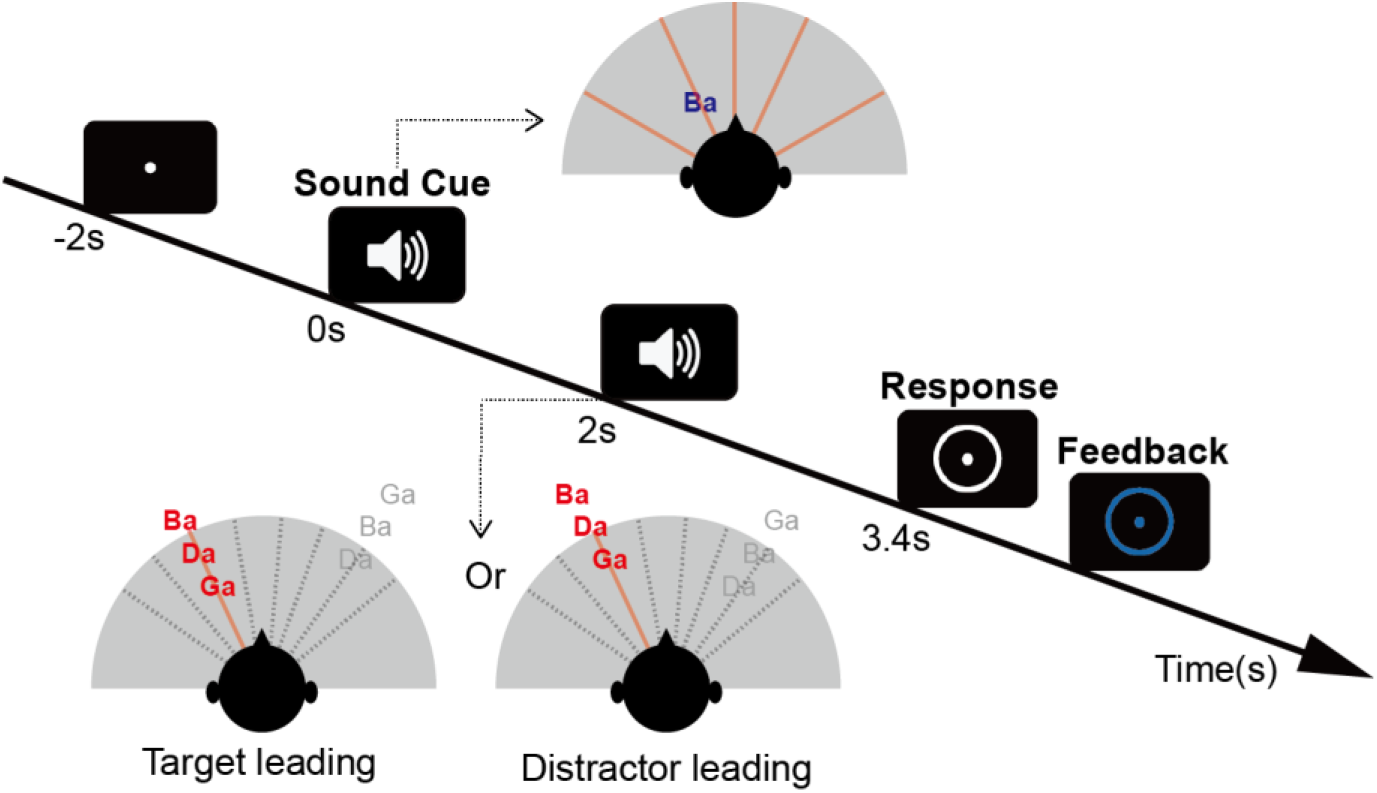
Trial design. A fixation dot appears at the center of the screen to instruct the listeners to fixate their gaze. An auditory cue of 400ms duration (the spoken syllable /ba/) begins 2 s later (at time zero) from one of five spatial locations chosen pseudo-randomly on each trial, indicating the location from which the target will be presented (in the top inset diagram, the /ba/ cue, in blue, is shown as coming from roughly 45 deg to the left, while the five potential target directions are shown by red radial lines). After a preparatory period (0 s - 2 s), two sound streams made up of random sequences of /ba/, /da/ & /ga/ (spoken by the same talker) are presented. The target stream (colored red) appears from the cued direction while a distractor stream (colored grey) appears from a different randomly chosen direction. In each trial, the stream beginning first is selected randomly, and the other stream begins 200 ms later (in the bottom inset diagrams, the target begins first for the left example, but second in the right example, while the distractor is presented from a location to the right; potential distractor locations are shown by gray dashed lines). After the two streams finish playing (3.4 s after the sound cue is presented), a white circle appears on the screen, indicating that it is time to report the target sequence. Immediately after the response is given, the circle changes color to provide feedback (blue to indicate a correct response or red to indicate an incorrect response).

A circle appeared around the fixation dot 3.4 s after the sound cue (after both target and distractor sequences finished playing), indicating the period during which subjects could record their responses. After entering their response, the fixation dot and response circle changed color to indicate whether the subject correctly reported the three target syllables (blue circle), or misidentified one or more syllables (red circle).

Participants performed 12 statistically identical blocks, each made up of 40 trials (for a total of 480 trials per subject). The order of the trials within each block was random, with the constraint that each of the five target locations was presented an equal number of times (each 8 times per block). Thus, over the course of the 12 blocks, each subject performed 96 trials with the same target location.

The syllables /ba/, /da/, & /ga/, spoken by the same female talker, were used both for both the auditory cue and to make up the target and distractor streams. The auditory cue was a single presentation of syllable (/ba/) with the spatial attributes of the upcoming target. The three-syllable target and distractor sequences consisted of random sequences of the syllables, chosen with replacement, and chosen independently for the target and distractor on each trial. All syllables were presented over headphones at a sound level of 70 dB SPL.

We varied the interaural time difference (ITD) of the stimuli to manipulate their perceived lateral position. Target sequences had ITDs of −600, −250, 0, 250, or 600 μs (roughly corresponding angular locations of −60°, −25°, 0°, 25°, 60°; Wightman and Kistler, 1992; Smith and Price, 2014).

On each trial, the distractor stream ITD was chosen to have an ITD that differed from the target ITD by one of 8 increments (−600, −450, −300, −150, 150, 300, 450, 600 μs), subject to the constraint that the absolute value of the resulting ITD value never equaled or exceeded the ethological range (max ITD magnitude of 700 μs; Feddersen et al., 1957; Kuhn, 1977). For example, if the target ITD was far to the right (target ITD: +600 μs), the distractor ITD was chosen from the set 0, 150, 300, or 450 μs (there were no possible ITDs farther to the right); if the target ITD was to the mid left (target ITD: −250 μs, as in Figure 1), then the distractor ITD was set to either −550 or −400 μs (to the left of the target) or −100, 50, 200, 350 or 500 μs (to the right of the target). This restriction was imposed to ensure that none of the trials was too easy, with very large separations between the target and the distractor.

### 2.3 Behavioral analysis

We calculated the percentage of correctly recalled syllables for each one of the three syllables in the target stream. For each of the syllables, we separately analyzed data from each of the 5 possible target locations, broken down based on whether the target or the distractor stream was temporally leading. Data were collapsed across the different distractor locations.

### 2.4 EEG analysis

#### 2.4.1 EEG data acquisition and preprocessing

EEG data was recorded with 64-channel Biosemi ActiveTwo system in an Eckel sound treated booth while participants performed the tasks. Two additional reference electrodes were placed on the mastoids. The stimulus timing was controlled by Matlab (Mathworks, Natick, MA) using the Psychtoolbox 3 extension (Brainard, 1997). EEG analyses included plotting scalp topographies using the EEGlab toolbox (Delorme and Makeig, 2004) and performing other functions in the Fieldtrip toolbox (Oostenveld et al., 2011).

EEG data from the correct trials were referenced against the average of the mastoid channels and down-sampled to 256 Hz. EEG data was then epoched from the sound cue onset to the end of the presentation period. Each epoch was baseline corrected by subtracting the mean from the baseline period (the 100 ms prior to the auditory cue). After baseline correction, trials with a maximum absolute value over 80 microvolts were rejected to remove artifacts (Delorme et al., 2007). Two subjects with excessive artifacts were removed from further EEG analysis (less than 60% trials remaining in at least one condition after artifact rejection). For the remaining 26 subjects, there were at least 92 trials remaining for each condition after artifact rejection. To equate the number of trials, 92 trials were randomly sampled for each condition for each subject for all subsequent analysis.

#### 2.4.2 Analysis of peak alpha power in frontocentral and parieto-occipital electrodes

For each epoch, the power spectrum was calculated over the 1s long period before the stimulus onset, thereby avoiding inclusion of any strong evoked activity. Data segments were zero padded to achieve a resolution of 0.1 Hz. For each subject and condition, the power spectra were averaged across trials (96 trials per condition) to estimate the spectrum for each EEG channel.

We were interested in whether peak alpha frequency varied systematically across the scalp. To assess this, we divided electrodes into frontal, frontocentral, and parieto-occipital groups based on their locations on the scalp (see Figure 3A). The across-trial average power spectra were then averaged across electrodes within each electrode group. The peak alpha frequency in each electrode group was found by determining the local maxima of the average power spectra within the 7-14 Hz band. If there were multiple peaks within this alpha range, the peak with the maximum height was selected.

For more than half of the subjects, there was no clear alpha peak in the frontal electrode group. Therefore, the frontal electrode group was excluded from this and any further alpha analysis. Similarly, any subject for whom the alpha peak could not be detected in at least one of the conditions for either frontocentral or parieto-occipital groups was excluded from further alpha analysis. One left-handed subject was excluded for this reason.

#### 2.4.3 Analysis of individualized parieto-occipital alpha frequency power

We wished to analyze how individual parieto-occipital alpha power changed with the spatial focus of auditory attention. For all subjects with identifiable peak alpha frequencies, we defined the individual parieto-occipital alpha frequency (IPAF) in the parieto-occipital electrode group. The IPAF was calculated by averaging the EEG power spectra across all trials in all conditions and across all parieto-occipital electrodes, then finding the peak frequency.

Once the IPAF was determined for each subject, we filtered all of their EEG data across the whole scalp with a 2 Hz wide bandpass FIR filter centered on the IPAF (IPAF ± 1Hz). We applied a Hilbert transform to the bandpass filtered data to extract the individualized alpha energy envelope, and took the magnitude of the transformed data. For each electrode and target location, we calculated the time course of the IPAF power for each trial and then averaged across trials to estimate the individualized induced alpha power time course (Snyder and Large, 2005). We baseline corrected the average IPAF power against 1s before the cue onset. The mean power averaged over the baseline period was subtracted from the IPAF power at each electrode. The resulting data was then divided by the standard deviation of the baseline period. We then calculated spatial z-scores of IAPF power for each electrode on the scalp by subtracting the mean IPAF power averaged across all electrodes and normalizing against the global field power (Murray et al., 2008; Skrandies, 1990). For each target location, we calculated the time course of the average IPAF power spatial z-scores.

To determine whether the direction of attention significantly altered the topographic distribution of alpha power, we contrasted the two extreme conditions: when subjects attended the leftmost target (target ITD: −600 μs) and when they attended the rightmost target (target ITD: +600 μs). Using the FieldTrip toolbox with Matlab, we performed a group-level analysis of GFP normalized IPAF power. For each subject, we computed the average of IPAF power over the whole trial for each electrode (0-3.4 s) for the leftmost and the rightmost conditions, resulting in two scalp topography plots for each subject. We then performed a spatial clustering analysis with FieldTrip to find clusters of electrodes across which the GFP normalized IPAF power differed significantly in these two topography plots. Multiple comparison was controlled by undertaking a Monte Carlo permutation test with 1000 random iterations using Fieldtrip. The cluster-based control has a type I error level of α = 0.05. The resulting clusters were then used to define the set of electrodes to combine for further analysis.

IPAF power time courses of all electrodes within each statistically significant cluster were averaged to produce one IPAF power time course for each cluster. A temporal clustering analysis was performed on the cluster-based time courses across the 5 target locations to test whether IPAF power changed significantly with direction of attention. Each time course was divided into 200 ms long time bins. A linear regression model was applied for each time bin (independent variable of target direction, dependent variable of magnitude of normalized IPAF power). An ANOVA was applied to the linear regression model and Bonferroni correction was performed to control for multiple comparisons. This analysis identified time windows exhibiting significant variation in IPAF power for different directions of attention.

## 3 Results

### 3.1 Behavior

Overall, participants were accurate in reporting the target sequence. All subjects were able to perform significantly above chance level (33%).

We conducted an ANOVA to examine how the percentage of correct responses varied across conditions. Given the sample size (N=28), we checked the normality of the sample distributions using the Lilliefors normality test and found that all distributions passed (*P*>0.05) before performing parametric statistical tests. The multi-way ANOVA had main factors of target location (five ITDs), leading stream (target or distractor), and syllable temporal position (first, second, or third). There was no main effect of target location [F_(4,839)_=1.14, *P*=0.34] or of leading stream [F_(1,839)_=2.41, *P*=0.12]. Furthermore, there was no significant interaction between target location and either of the other factors [target location × leading stream: F_(4,839)_=1.22, *P*=0.30; target location × syllable position: F_(8,839)_=0.22, *P*=0.99], indicating that task performance did not vary with target location. However, there was a significant main effect of syllable position [F_(2,839)_=11.1, *P*<0.001] and a significant interaction between syllable position and leading stream [F_(2,839)_=63.9, P<0.001].

Figure 2 shows the percentage of correctly recalling each syllable in the target leading (left panel) and distractor leading (right panel) conditions (collapsing across target location). The data show that when the target leads, performance decreases from the first target syllable to the last target syllable, while this pattern reverses when the target lags.

**Figure 2.**
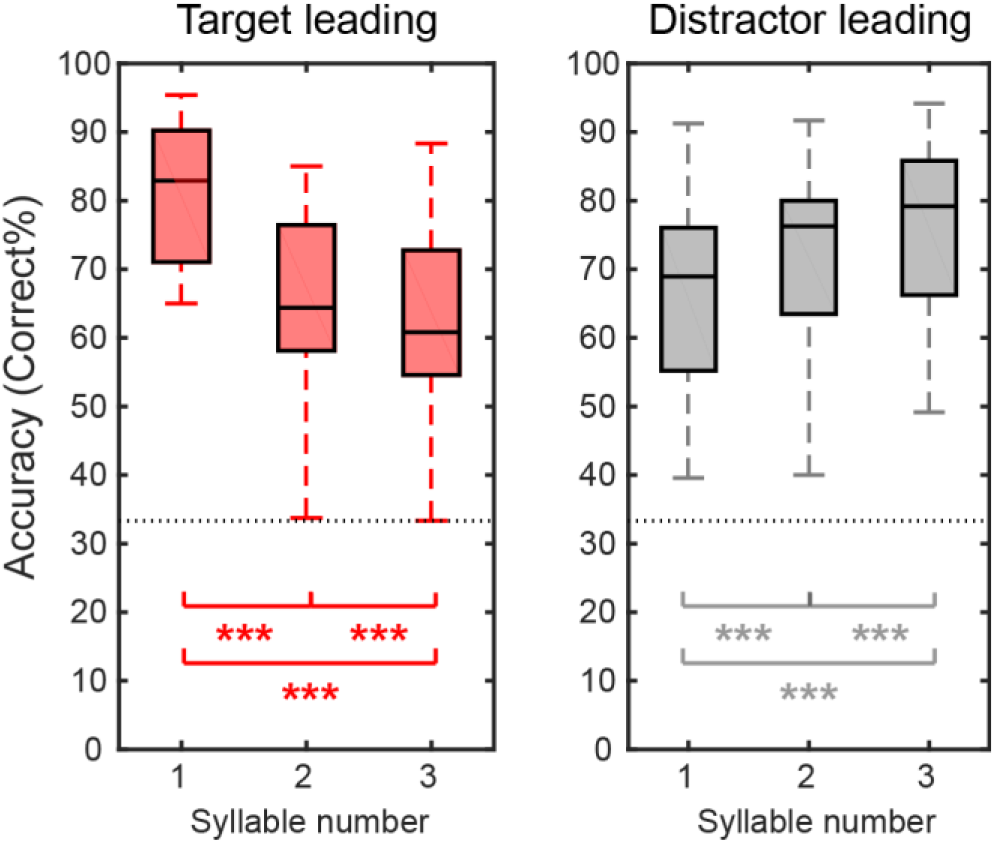
Behavioral performance averaged across subjects for the target leading (left panel) and distractor leading (right panel) conditions. Within each panel, we show boxplots of percent correct responses (central black line shows the across-subject mean; boxes show the 25^th^ – 75^th^ percentile ranges; error bars show the range from minimum to maximum) for each syllable position within the target stream. The horizontal gray dashed line shows chance performance (33%). Asterisks indicate percentages that differ significantly from one another based on post hoc tests (* *p*<0.05; ** *p*<0.01; *** *p*<0.001).

To test the significance of these observations, we did post-hoc tests. We corrected for multiple comparisons by calculating post-hoc test statistics using the Benjamini-Hochberg FDR correction (Benjamini and Hochberg, 1995). Our post hoc tests (paired t-tests) showed that the percentage of correct response decreases systematically with syllable number when the target stream leads (syllables 1 vs. 2, t_(27)_=9.83, *P*<0.001; syllables 1 vs. 3, t_(27)_=9.49, *P*<0.001; syllables 2 vs. 3, t_(27)_=3.78, *P*<0.001), but performance improves from syllable to syllable when the distractor stream leads; (syllables 1 vs. 2, t_(27)_=4.94, *P*<0.001; syllables 1 vs. 3, t_(27)_=6.99, *P*<0.001; syllables 2 vs. 3, t_(27)_=3.80, *P*<0.001).

We tested whether handedness influenced behavioral performance using a two-sample t-test at the group level. We found no significant difference between left-handed and right-handed groups’ performance [t_(14)_=0.53, *P*=0.60].

### 3.2 Peak alpha power in frontocentral and parieto-occipital electrodes

For each subject and target location, we computed the frequency that had the greatest average power within the alpha range separately for frontocentral electrodes and parieto-occipital electrodes. These results, shown in Figure 3, suggest that alpha peak frequency is higher in parieto-occipital electrodes than in frontocentral electrodes.

**Figure 3.**
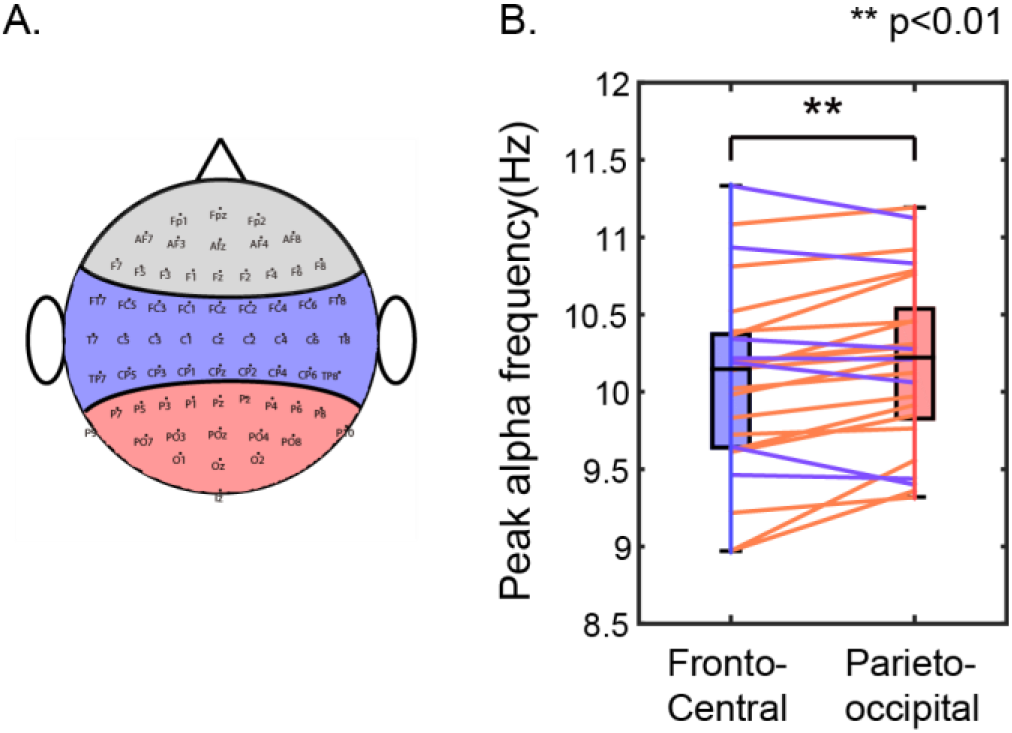
Peak alpha power is greater in parieto-occipital electrodes compared to frontocentral electrodes. A. Definition of electrode groups. Blue and red area represents the electrodes included in frontocentral and parieto-occipital groups; frontal electrodes (in gray) showed less consistent and robust alpha, and therefore were not analyzed. B. Comparison of peak alpha frequency in the frontocentral and parieto-occipital electrodes. Box plots show group comparisons (as in Figure 2). Blue and red colored dots plot results from individual subjects; colored lines connect peak frequencies in the two electrode groups for a given subject, with the color of the connecting line shows which peak frequency is greater: blue lines indicate higher peak frequency in frontocentral electrodes, while red lines indicate higher peak frequency in parieto-occipital electrodes.

We explored the statistical significance of these observations by conducting a multi-way repeated-measure ANOVA on peak alpha frequency with main factors of electrode group (frontocentral and parieto-occipital) and target location (five ITD values). There was no significant main effect of target location on alpha peak frequency [F_(4,249)_=1.63, *P*=0.12)] and no significant interaction between target location and electrode group [F_(4,249)_=0.80, *P*=0.53]. However, there were significant main effects of both subject identity [F_(24,249)_=220, *P*<0.001] and electrode group [F_(1,249)_=59.27, *P*<0.001], as well as significant interactions of subject identity with both electrode group [F_(24,249)_=6.61, *P*<0.001] and target location [F_(96,249)_=1.51, *P*=0.022]. These results suggest that while peak alpha frequency varies across subjects, there is a consistent difference in the peak alpha frequency in frontocentral and parieto-occipital electrodes. Post-hoc t-test reveals that the alpha peak frequency is generally higher in the parieto-occipital region than in the frontocentral region [t_(24)_=2.99, *P*=0.006; paired t-test; Figure 3B].

We also conducted a group level two-sample t-test to examine whether alpha peak frequency varied with handedness. We did not find any significant difference in peak alpha frequency between the left-handed and the right-handed groups [t_(23)_=1.37, *P*=0.18].

### 3.3 Individualized parieto-occipital alpha frequency power

We explored how the topographic distribution of individual parieto-occipital alpha frequency power changed with the spatial focus of auditory attention. Figure 4A shows a scalp topography plot of the difference, at each electrode, between the average IPAF power when attending far left (target ITD: −600 μs) and far right (target ITD: 600 μs). Significant electrodes from spatial clustering results are overlaid on the topography plot. A positive cluster was found on the left parieto-occipital region (*P*=0.007) and a negative cluster was found on the right hemisphere (*P*=0.009). This result is in consistency with the previous literature, which reports that alpha oscillation power decreases in the hemisphere contralateral to an attended location and increases in the hemisphere ipsilateral to an attended location (Banerjee et al., 2011; Kelly et al., 2006; Klimesch, 2012; Worden et al., 2000).

**Figure 4.**
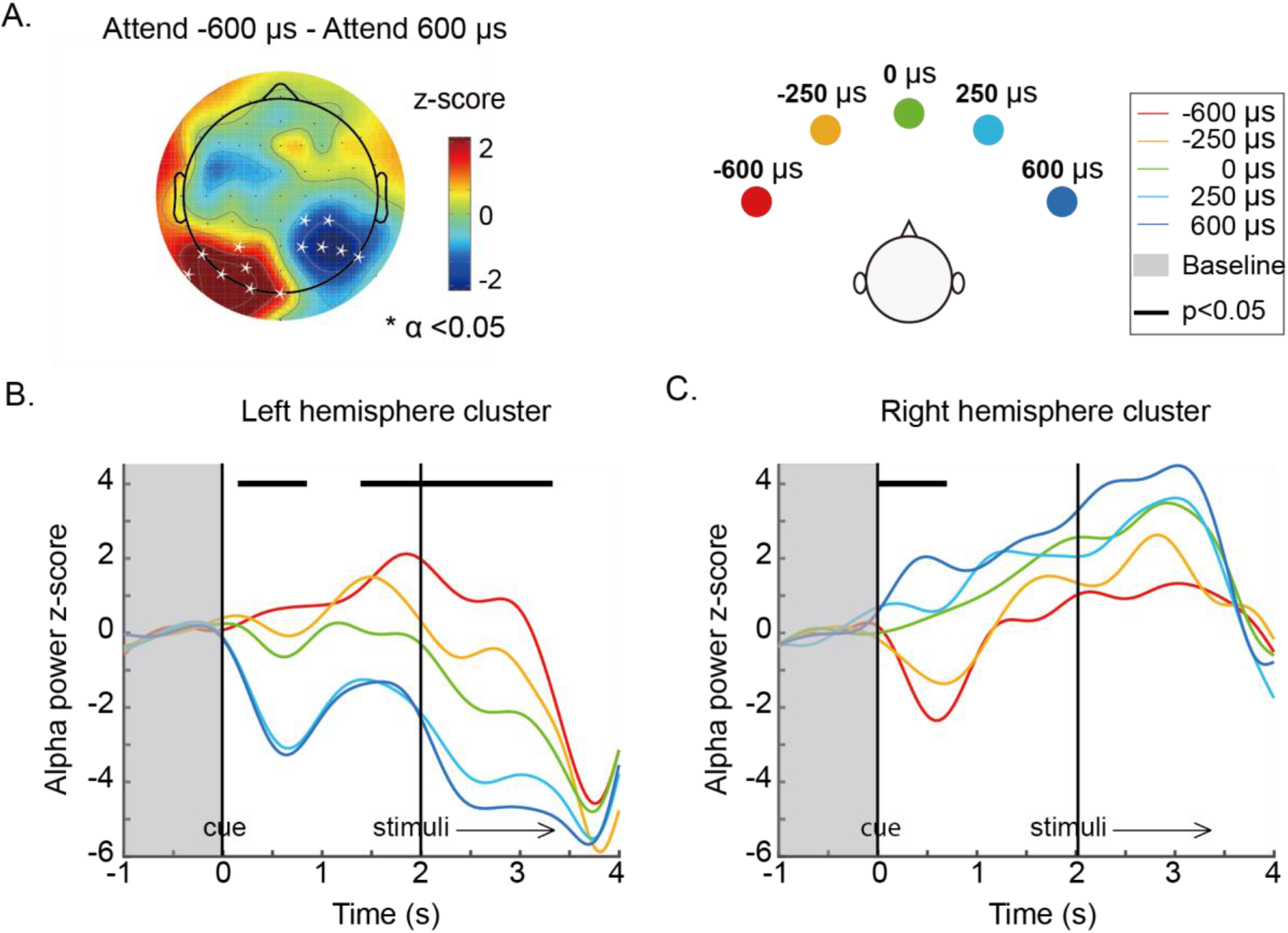
A. Topography of alpha power z-score difference between attend far left (target ITD: −600 s) and attend far right (target ITD: 600 s) averaged over the whole trial (0-3.4s). Channels within a cluster in which alpha power changes significantly with direction of attention are marked by asterisks. B & C. Time courses of alpha power dynamics across the trial period for the left parieto-occipital cluster and the right parieto-occipital cluster. Individualized parieto-occipital alpha power was averaged within the left and right clusters shown in A and plotted as a function of time for each target location. Data were divided into 200 ms long time bins and averaged within each time bin. Asterisks at the top of the plot (forming continuous bars, visually) show the time windows in which there was a statistically significant effect of target direction on alpha power after Bonferroni correction. Vertical black lines illustrate the onsets of auditory cue and stimuli. Shadowed areas represent the baseline period.

To investigate the influence of handedness on alpha lateralization, we calculated the degree of alpha lateralization for each subject in each attention condition. Specifically, for each target direction, we subtracted the IPAF power averaged across the right hemisphere cluster (which is generally negative for attention to the left and positive for attention to the right) from the IPAF power averaged across the left hemisphere cluster (which is generally positive for attention to the left and negative for attention to the right). ANOVA revealed a significant main effect of target location on lateralization (F_(4,129)_=7.33, *P*<0.001) but no significant effect of handedness (F_(2,129)_=2.68, *P*=0.10) and no significant interaction between handedness and attention focus (F_(4,129)_=0.08, *P*=0.99).

Within each cluster, the average was taken to render the time courses of IPAF power in each attention focus condition (Figure 4B & 4C). The IPAF power time courses reveal that alpha power increases systematically as the focus of spatial attention shifts from left to right in the left parieto-occipital cluster, and decreases systematically in the right parieto-occipital cluster.

To test for the significance of these effects, we performed a linear regression analysis for each 200 ms long time bin. Bonferroni corrected results revealed that alpha power varies significantly with direction of attention in a number of time periods. In the left parieto-occipital cluster, alpha power changed significantly with target direction immediately after the cue (0.2-0.8s) and again from right before the target/distractor sound began until it finished played (1.6-3.2s; Figure 4B). The opposite trend is seen throughout the trial in the right parieto-occipital cluster; however, this variation was only statistically significant immediately after the cue appeared (0-0.8s; Figure 4C).

Given that both left and right parieto-occipital clusters show significant differences in alpha power during the preparatory period, we undertook a final analysis to quantify how alpha power varied across the scalp with direction of attentional focus. To this end, for each electrode we averaged alpha power in the preparation period (0s – 2s, after the onset of the cue for where to attend, but before the onset of the target and distractor stimuli) for each target location.

Figure 5A shows these average alpha power values across the scalp. As attentional focus shifts from left to right, there is a clear change in the power of alpha in parieto-occipital electrodes: alpha decreases systematically in the left parieto-occipital electrodes and increases systematically in the right parieto-occipital electrodes. To visualize these changes, we took the average alpha power over the left and right parieto-occipital clusters identified above (see Figure 4A), and plotted the mean activity as a function of target ITD (Figure 5B). These results show that in left parieto-occipital electrodes, alpha power is significantly greater than baseline when attention is directed ipsilaterally (far left of the left panel of Figure 5B); decreases as attention shifts to contralateral, right exocentric space (moving from left to right in the panel); and is significantly below baseline (reduced alpha) when attention is focused on the right. Consistent with past work on parieto-occipital responses during attention, our results are not symmetric (Haegens et al., 2011; Ikkai et al., 2016). In the right parieto-occipital cluster, alpha power is greater than baseline when attention is directed ipsilaterally (far right of the right panel of Figure 5B); decreases as attention shifts to contralateral, left exocentric space (moving from right to left in the panel); but never falls significantly below baseline, even when attention is focused on the far left.

**Figure 5.**
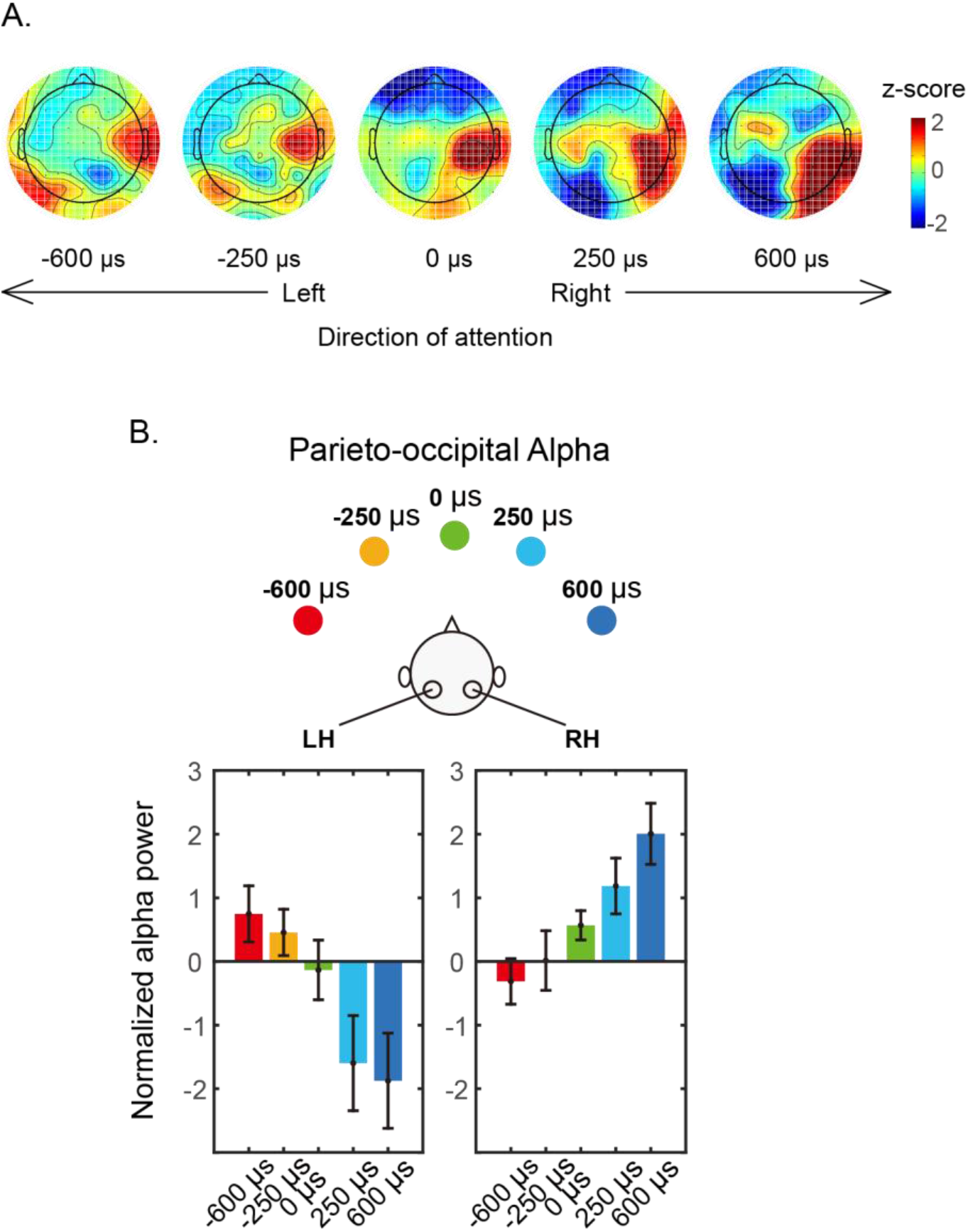
Parieto-occipital alpha power during the attentional preparatory period (−2 to 0s). A. Topographies of alpha power z-scores for the five different target locations. B. Parieto-occipital alpha averaged within the left and right parieto-occipital clusters revealed by clustering analysis.

## 4 Discussion

We tested subjects engaged in a challenging auditory spatial attention task and observed how alpha oscillation power changed during task performance. Importantly, the task was designed to require sustained auditory spatial attention: two competing streams, each a sequence of three syllables selected from the same set of tokens, spoken by the same talker, and overlapping in their time of presentation, were presented with two different ITDs. Subjects were asked to report the target sequence from the direction indicated by an auditory cue at the start of each trial, while ignoring the distractor (from an unknown direction). In order to force listeners to rely on spatial cues, which of the streams began first was random from trial to trial, and counter balanced.

Behaviorally, listeners were good on the task. Still, the pattern of behavioral results depended on the exact stimulus configuration. Specifically, when the target began before the distractor, listeners were very good at reporting the first target syllable; however, they were worse at reporting the second, and even worse at reporting the third. Given that the distractor began playing before the second syllable, this decrease in performance with syllable number is not very surprising, and likely reflects a combination of both energetic masking (e.g., see Arbogast et al., 2002; Brungart, 2001) and more central, cognitive interference (e.g., see Shinn-Cunningham, 2008). The opposite pattern occurs when the distractor began first: performance was worst for the first syllable, better for the middle syllable, and best for the final syllable. Again, this pattern makes sense. The sudden onset of the distractor at the start of the presentation undoubtedly grabs attention involuntarily (see, for example, Buschman and Miller, 2007; Conway et al., 2001; Elhilali et al., 2009). The first target syllable begins only 200 ms after the distractor; this brief delay is close to the limit for how quickly listeners can shift attention away from the salient distractor onset, which impacts the ability to report the first target syllable. Over time, spatial attention to the correct stream builds up, leading to better focus as the presentation continues, consistent with some previous studies of auditory attention (Best et al., 2008; Dai et al., 2018). Moreover, the distractor sequence ends before the third target syllable begins. Together, these effects lead to improvement in performance from syllable to syllable on trials where the distractor begins first. Overall, however, it is clear that listeners were able to perform the task well, and that they relied on auditory spatial cues to perform the task.

### 4.1 There are significant subject differences in alpha peak frequency

We observed that alpha peak frequency varied across individuals, consistent with previous reports (Basar, 2012; Bodenmann et al., 2009; Klimesch, 1999). For instance, we found that during the task, the peak alpha frequency in individual listeners’ parieto-occipital electrodes ranged from 9-11.3 Hz (standard deviation of 0.6 Hz). This observation argues for the importance of analyzing alpha in subject-specific ways (Haegens et al., 2014). We therefore estimated the alpha peak frequency for each subject and used this to estimate alpha power in all subsequent analysis. Using subject-specific analysis of alpha ensures that we get the cleanest, most robust measures of how alpha power changes with task demands.

### 4.2 Multiple generators of alpha are engaged during selective auditory spatial attention

We separately analyzed the dominant frequency of alpha power in frontal, frontocentral, and parieto-occipital electrodes. While there was not a robust peak in alpha power in frontal electrodes, we found clear peaks in frontocentral and parieto-occipital sensors. Moreover, we found consistent differences in the frequency of the dominant alpha peak in frontocentral versus parieto-occipital electrodes. Specifically, the peak frequency of alpha in the frontocentral electrodes is significantly lower than in the parieto-occipital electrodes: 19 out of 25 subjects showed peak frequencies of frontocentral alpha that were lower than for parieto-occipital alpha (see Figure 3). This difference in alpha peak frequency provides strong evidence for multiple generators of alpha activity during auditory tasks, leading to different scalp topographies (one stronger over frontocentral electrodes and one stronger over parieto-occipital electrodes).

A number of previous studies have reported changes in alpha oscillation power during challenging auditory tasks. However, different studies attribute different roles to these oscillations and what they signify. For instance, previous studies have reported that during auditory spatial attention, alpha activity tends to lateralize, increasing ipsilateral to the direction of attention and decreasing contralateral to the direction of attention in both temporal (e.g., Hartmann et al., 2012) and posterior brain regions (e.g., Banerjee et al., 2011; Wöstmann et al., 2016). Increases in alpha power have been associated with increases in cognitive load in temporal as well as posterior portions of the brain (Van Dijk et al., 2010; Wilsch and Obleser, 2016; Woestmann et al., 2017), and alpha power increases with listening effort (Woestmann et al., 2015). Prestimulus alpha has been shown to reflect decision processes (Woestmann et al., 2019). The current study is consistent with the various reports of alpha power reflecting a range different functions during auditory task performance; our results suggest that during our auditory spatial attention there are multiple generators of alpha, which come from different neural regions and thus reflect different cognitive processes (see also Weisz et al., 2014). Indeed, the point that multiple alpha generators likely contribute during different auditory tasks has been put forth in a recent review paper (Strauß et al., 2014).

Some previous studies have shown that in addition to affecting alpha power, task engagement and even task load can influence peak alpha frequency (e.g., Basar, 2012; Haegens et al., 2014). In the current task, even though we expected to see (and saw) changes in parietal alpha power with the direction of attention (see the discussion below), we did not find any changes in alpha peak frequency when we varied target location. This result makes sense, given that task performance (and thus task difficulty) was similar for different target locations. Our results are consistent with the idea that, regardless of the specific direction of attention (target location), the same brain networks are engaged in performing the same basic cognitive functions during the task, which leads to the same frequencies of alpha oscillations across all conditions.

Our study methods limit our ability to localize the generators of observed neural activity (EEG measures with a small number of sensors and without any subject-specific models of anatomical structure); thus, we cannot, from the current results, make strong claims of where the different neural generators of alpha oscillations lie. Given how EEG signals propagate to the scalp, parietal sources of alpha are likely to dominate the observed responses from parieto-occipital electrodes, while sources more frontal sources likely dominate the responses in frontocentral electrodes. Regardless, our results provide good evidence that there are at least two different generators of alpha oscillations during our auditory task.

The alpha power we observe in frontocentral electrodes central alpha range oscillation could be a mu rhythm, related to motor planning (Llanos et al., 2013; Sabate et al., 2012). The frequency range of mu rhythms (7.5-12.5Hz) overlaps with alpha. In our task, the task-related modulation of frontocentral alpha led to greater alpha power in right-hemisphere electrodes, but not in left-hemisphere electrodes, and did not vary significantly with the direction of attention (consider Figures 4A & 5A). This pattern is consistent with a right-handed motor response during the task, which leads to an increase of mu oscillations over right motor cortex [related to suppression of movement of the left hand; (Pfurtscheller et al., 2006, 2000; Wolpaw et al., 2002)]. While we tested both right- and left-handed subjects, even left-handed subjects used the numeric keypad (on the right side of a keyboard) to enter their responses, consistent with the observed results. Alternatively, the more frontal alpha source could be from auditory sensory cortex, which has been reported to generate alpha power that fluctuates during auditory task performance (Frey et al., 2014).

Future work is needed to tease apart how different neural generators behave during tasks like that used here. To address these questions, neuroimaging techniques with better spatial resolution should be employed to allow localization of the underlying neural generators.

### 4.3 Parieto-occipital alpha power topography reflects the lateral position of auditory spatial attentional focus

We found that when auditory attention is covertly oriented to a particular spatial location, alpha power in parieto-occipital electrodes lateralizes, increasing in the electrodes ipsilateral to the direction of attention and decreasing in the contralateral electrodes. These results are consistent with previous studies of both visual and auditory spatial attention (Frey et al., 2014; Sauseng et al., 2005; Strauß et al., 2014; Thut et al., 2006; Wöstmann et al., 2016). Our study extends these previous findings by contrasting the topographic distribution of alpha power as a function of the lateralization of attention, testing five different target directions ranging from far left to far right: the farther lateralized the focus of attention focus, the greater the lateralization of parietal alpha power. While previous studies have shown that it is possible to decode the focus of visual attention from the distribution of alpha power (Foster et al., 2016; O’Sullivan et al., 2015; Rihs et al., 2007; Samaha et al., 2017), the topographic distribution of alpha power has, to our knowledge, not been shown previously to change systematically with the direction of auditory spatial attention. Previous studies of auditory spatial attention have generally only considered how alpha is distributed during attention to one location on the left versus attention to a symmetric location on the right, or even for dichotic sounds (a sound presented only to the left ear and a different sound presented only to the right ear).

The pattern of alpha lateralization that we report is consistent with the theory that alpha reflects a suppressive or inhibitory mechanism (Jensen and Mazaheri, 2010; Klimesch, 2012; Klimesch et al., 2007; Strauß et al., 2014). Specifically, parietal cortex has a contralateral bias, primarily encoding information from contralateral exocentric space (Kaiser et al., 2000; Schonwiesner et al., 2006; Teshiba et al., 2013). Given that spatial auditory processing engages retinotopically organized parietal maps of contralateral space (Huang et al., 2014), increases in alpha power ipsilateral to the direction of attention are consistent with suppressing interfering information about events in the opposite direction. Attention to a particular retinotopic location is likely to cause an alpha-linked suppression of information in subnetworks of the brain representing other retinotopic locations. Our observation of a gradation of parietal alpha power lateralization that reflects the exact attentional focus is consistent with the theory that local alpha power modulation “reflects changes in the excitability of populations of neurons whose receptive fields match the locus of attention” (Ikkai et al., 2016; Klimesch, 2012).

Visual attention studies show that the topography of parietal alpha varies not only with left-right lateral angle, but also with elevation. Although perception of auditory elevation is substantially less precise than perception of auditory lateral angle (which is already much less precise than visual perception of angle), it would be interesting to explore whether changes in the elevation of auditory spatial attention (e.g., using free-field speakers to provide rich, realistic auditory elevation cues) also affect the distribution of parietal alpha power.

### 4.4 Is the frontoparietal network truly a supramodal spatial attention network?

As discussed above, we find that just like in both vision (e.g., Kelly et al., 2006; Worden et al., 2000) and touch (e.g., S. Haegens et al., 2011), spatial attention directed to an auditory target causes a shift in parietal alpha power (with relatively greater power in parieto-occipital electrodes ipsilateral to the direction of attention; Banerjee et al., 2011; Wöstmann et al., 2016). Given the difficult in localizing the sources of the observed EEG results, however, this alone provides relatively weak support for the idea that the spatial attention network is shared between vision and audition.

A couple of neuroelectric studies have directly contrasted parietal alpha lateralization for visual and auditory spatial attention, and found clear differences in topography (Frey et al., 2014; Banerjee et al., 2011). Indeed, in MEG, which provides better resolution of deeper brain structures than does EEG (Frey et al., 2014), auditory spatial attention was shown to modulate the lateralization of alpha in auditory sensory cortex, but not visual spatial attention. Yet, although there were differences in alpha topography for visual and auditory spatial attention tasks, alpha lateralization in parieto-occipital regions was similar across modalities. Thus, while there are multiple generators of alpha during auditory tasks, the alpha associated with suppression in parietal cortex may well reflect the same cognitive mechanism during visual and auditory spatial attention.

Recent fMRI work that uses subject-specific definitions of regions of interest (ROIs) based on functional localizers supports the view that auditory spatial processing engages the very same regions that are always engaged during visual processing. Specifically, distinct frontal executive control regions that are biased towards visual processing are differentially more engaged during auditory attention and working memory tasks (Kong et al., 2014; Michalka et al., 2016; Noyce et al., 2017). Importantly, these executive regions, together with retinotopically mapped regions in parietal corte, form a coherent network that is seen during fMRI resting state in both subject-specific ROI analysis (Michalka et al., 2016) and that emerge at a group level from the large-scale connectome dataset (Tobyne et al., 2018). These results lend further support to the view that the frontoparietal visual spatial attention network is also engaged during auditory spatial processing.

Another piece of evidence for the supramodal nature of parietal representations is the common asymmetry seen in the information representation across modalities. The right hemisphere dominance theory posits that the left hemisphere represents information from right exocentric space, whereas the right hemisphere, while biased towards representing left exocentric space, also represents ipsilateral information (Huang et al., 2014; Mesulam, 1999; Okazaki et al., 2015; Pouget and Driver, 2000; Shulman et al., 2010). This asymmetry helps explain why hemifield neglect is common for sources in left exocentric space (i.e., in patients with right lesions in parietal cortex that destroys the only information about leftward sources), but uncommon for right exocentric space (Heilman and Abell, 1980). This kind of left-right asymmetry is seen not only in past results, but in our current auditory spatial attention data.

Specifically, previous neuroelectric studies in both vision (e.g., Ikkai et al., 2016) and touch (e.g., Haegens et al., 2011) report greater modulation of parietal alpha power when attention is directed to left compared to right exocentric space. We see the same asymmetry. During the preparatory period (following the cue but before the stimuli began), alpha power in the left electrode cluster decreased below baseline when attention was focused on the right, and increased above baseline when attention was focused on the left. In contrast, in right electrodes, preparatory alpha power never decreased significantly below baseline, even when attention was directed to the far left. Furthermore, attentional modulation of parietal alpha was significant throughout the presentation of the target-distractor stimuli in left parieto-occipital electrodes, but less robust (and not statistically significant) during the stimuli in right parieto-occipital electrodes.

We included equal numbers of left- and right-handed subjects in the study with the intention of studying effects of atypical hemispheric asymmetry in left-handed subjects. However, we did not find any significant difference between left- and right-handed subjects in any of our analyses. For this reason, we collapsed all of our data across these groups in the presented results. Given the number of subjects we were able to test, combined with the relatively low incidence of atypical hemispheric dominance even in left-handed participants (Knecht, 2000), this failure to find an effect is not particularly surprising. Future studies with a prescreening procedure to test for hemispheric dominance, and separating participants into groups based on this independent measure, would undoubtedly shed more light on how parietal processing is affected when subjects have an atypical spatial representation (e.g., Cai et al., 2013).

## 5 Conclusions

We studied how individualized parietal alpha power shifts as a function of the lateral direction of auditory spatial attention. We presented auditory targets from one of five azimuth locations by varying ITD from −600 μs to +600 μs. We found unique alpha peak frequencies over frontocentral and parieto-occipital electrodes, revealing the presence of at least two distinct generators of alpha oscillations during our task. The parieto-occipital alpha power was modulated by the lateral focus of attention, varying systematically with the focus of auditory attention. The similarity to previous results from other sensory modalities in alpha power lateralization, down to an asymmetry between alpha power changes in left versus right hemisphere, supports the view that the same cognitive processes are engaged during spatial attention across sensory modalities. Past fMRI evidence that the exact brain regions engaged by auditory spatial processing are part of the well-studied frontoparietal visual processing network; together with current results, the current study supports the idea that there is a common, supramodal spatial attention network.

## Acknowledgements

This work was supported by the National Institutes of Health [NIDCD R01 DC013825].

## Notes

The authors declare no financial conflicts of interest.

